# Evaluation of solubility and histological effect of 11-keto-β-Boswellic acid on the diabetic mice liver using FTIR-PCA analysis and microscopy

**DOI:** 10.1101/2021.01.27.428445

**Authors:** I.S. Al-Amri, F. Mabood, I.T. Kadim, A.Y. Alkindi, A. Al-Harrasi, S. Al-Hashmi, G. Abbas, B. Ahmed, J. Al-Shuhaimi, S.K. Khalaf, J. Shaikh

## Abstract

This study was designed to develop a rapid, sensitive, accurate, and inexpensive Fourier Transform Infrared Reflectance (FT-IR) Spectroscopy coupled with Principle Component Analysis (PCA) as a detection technique to evaluate the solubility of 11-Keto-β-Boswellic acid (KBA), from the gum resin extracted from the Omani frankincense, (*Boswellia sacra*) in the liver of STZ induced diabetic mice. This study also investigated the effect of KBA on the histological changes of hepatocytes of diabetic mice. Liver tissue samples from three groups of mice included normal control group, diabetic control group and diabetic group treated IP with KBA were scanned with FT-IR spectrophotometer in the reflection mode. FT-IR Spectra were collected in the wavenumber range from 400 to 4000cm^-1^ using ATR accessorry. The results of FT-IR Spectra were analyzed by using multivariate method Principle Component Analysis. The PCA score plot is an exploratory multivariate method indicated that there was a complete segregation among the three groups of liver samples based on change in variation of position of wavenumber in FT-IR spectra, which revealed that there is a clear effect of KBA solubility on treatments. The histological features showed an improvement in the liver tissues with normal structures of hepatocytes with exhibiting mild vacuolations in their cytoplasm. In conclusion, reflectance FT-IR spectroscopy coupled with PCA could be deployed as a new detection method for rapid, low cost and non-destructive method for evaluating of treatment effects in diseased liver tissue based on the solubility of KBA. Histological findings demonstrated the protective effective of KBA on improving the morphology of liver tissue in diabetic mice which resulted in complete recovery to the damage observed in diabetic control group.

**Summary Statement:** Reflectance FT-IR spectroscopy coupled with PCA has been deployed as a new rapid, inexpensive and non-destructive detection method to examine the solubility of 11-keto-β-Boswellic acid (KBA) in streptozotocin (STZ) induced-diabetes mice liver tissue following intraperitoneal treatment. Moreover, microscopic study of liver tissue histopathology revealed that KBA has a protecting effect against STZ damage.

## INTRODUCTION

Diabetes mellitus (DM) is a chronic metabolic disease categorized by hyperglycemia caused by insufficient insulin actions (Baynes, 2015). In 2020, the International Diabetic Federation (IDF) reported 463 million cases with diabetes worldwide, including 55 million cases in the Middle East and North Africa (MENA) region and 291 thousand in Oman. In 2017, 94.921 cases of diabetes were reported in Oman (Ministry of Health). Statistically, it has been estimated that the number of diabetes cases in MENA region will rise to 108 million by 2045 (IDF, 2020). DM causes several diseases such as neurological disorders, coronary artery disease, renal failure, cerebrovascular disease, limb amputation, blindness, and death (Middha et al., 2014; Shaikh and Shrivastava, 2014). Diabetes Type 1 and 2, gestational diabetes, impaired glucose tolerance and impaired fasting glycaemia are considered as the main types of diabetes (Baynes, 2015). Diabetes Type 1 is an autoimmune disorder whereas Type 2 is a metabolic disorder (Al-Haddad et al., 2016).

Human liver is the largest internal organ, with multifunction’s including regulation of glucose concentration in physiological and pathological conditions and prevention of excessive fluctuation (Cotoi and Quaglia, 2016). Hyperglycemia affects the metabolism of protein, carbohydrates, and lipids, leading to non-alcoholic fatty liver disease (NAFLD) (Mohamed et al., 2016). The condition of NAFLD can further develop to non-alcoholic steatohepatitis, cirrhosis, and ultimately hepatocellular carcinomas. Reports indicated that NAFLD affects up to 70–80% of people with type 2 diabetes mellitus and up to 30–40% of people with type 1 diabetes mellitus (Targher et al., 2018).

Medicinal components extracted from frankincense (*Boswellia* species) plant have been documented to possess health benefits and pharmaceutical properties such as anti-microbial anti-inflammatory, anti-cancer, anti-diabetic, antioxidant, and other analgesic activity (Al-Yasiry and Kiczorowska, 2016; Sabra and Al-Masoudi, 2014; Ni et al., 2012; Ammon, 2019; Al-Harrasi et al., 2013, 2014). The active derivatives of the plant are the boswellic acids found in the gum resin of *Boswellia* species with pharmacologically active pentacyclic triterpene molecules including 11-keto-β-boswellic acids (KBA) (Sabra and Al-Masoudi, 2014; Al-Harrasi et al., 2013). These active derivatives have been used to treat a number of inflammatory diseases, such as osteoarthritis, chronic colitis, bronchial asthma, pancreatic cells and tumors, human breast cancer cells (Saraswati and Agrawal, 2012), and hepatocellular carcinomas (Ahmed Hanaa et al., 2015).

Frankincense reported to significantly increse wound contraction rate following administration of KBA for 16 days on diabetic mice (Pengzong, et al., 2019). In another study, injected diabetic mice with KBA for seven weeks triggered infiltration of large number of lymphocytes to the pancreatic islands and the appearance of semi-isolated apoptosis cells was observed (Shehata et al., 2017). A decrease in the blood sugar level of a group of diabetic mice following injection with KBA has also been reported (Shehata et al., 2015). Pharmacokinetic studies, however, have evidenced low absorption of KBA in humans and rodents as these molecules have poor water solubility and strong tendency to self-aggregate (Wang et al., 2014; Hüsch et al., 2013; Büchele and Simmet, 2003; Sharma Sharma et al., 2004). The concentration of KBA declined with elimination of half-life after six hours of oral administration and suggested that medication is required after every six hours (Sharma et al., 2004). Therefore, strategies to improve its bioavailability need to be established, to ensure effective anti-inflammatory activity of KBA.

Fourier Transform Infrared Reflectance (FT-IR) is an effective tool used to determine the chemical characteristics of various sample forms including solid, liquid, or gases. FT-IR presents unique information by addressing cofactors, amino acids, and water molecules properties, with high structural parameters and interactions (Berthomieu and Hienerwadel, 2009. Advantages of this technique include rapid sample preparation and processing in solid forms for scanning with the FT-IR spectrophotometer followed by applying multivariate PCA method to build PCA model for exploring similarities and differences among different samples based on the solubility of the drug compound. This study was designed to provide analysis on the solubility of KBA from the gum resin extracted from the Omani frankincense (*Boswellia sacra*) in the liver of streptozotocin (STZ) induced diabetic mice and evaluate the effects of KBA on the morphological changes of liver tissue under disease conditions.

## MATERIALS AND METHODS

### Experimental design and tissue samples preparation

In present study, 18 female CD1 mice weighing 25–30 g and 10-12 weeks old were used. Initially, mice were divided into two groups: normal healthy control group (6 mice) and streptozotocin-induced group (12 mice). Six mice were kept in cages and exposed to a 12-h light/dark cycle in controlled room temperature and humidity. All mice had access to food and water *ad libitum*. One week after adaptation, the mice in control group (group 1) were injected with citrate buffer while the mice in group two were injected intraperitoneal (IP) with streptozotocin (180 mg/Kg) body weight in 10 mM citrate buffer (pH 4.5). Two weeks following injection with streptozotocin, blood samples were collected from the mice tail vein. Streptozotocin induced mice with blood glucose higher than 300 mg/dL were classified as diabetic mice and divided into two experimental groups each containing 6 mice. Diabetic control mice (group 2) received water only while group 3, diabetic mice were treated IP with KBA dissolved in water at doses of 25 mg/kg/day. Four weeks later after the final treatment, mice were anesthetized and sacrificed. Liver tissues for light microscopy screening were dissected and immersed in 10% buffered formaldehyde solution. For FT-IR analysis, liver tissues were frozen at −80°C in liquid nitrogen.

### FT-IR spectroscopy

Each tissue sample was scanned by FT-IR spectrophotometer to produce reflectance spectra at five positions of mice dried liver tissue samples with sixteen scans at each position to produce 80 spectra per sample. The averages of these 80 spectra per sample were used for building the multivariate PCA models.

### Principal Component Analysis (PCA)

The Unscrambler software X10.3 (CAMO Software, Oslo, Norway) and Microsoft Excel 2016 software were used to conduct the PCA models of the FT-IR spectral data. PCA was performed to determine the classification and segregation of the three groups of mice dried liver tissue samples based on the solubility of KBA. PCA models were built using the SVD algorithm. Briefly, PCA is a non-supervised data exploration multivariate chemometric method used to show the hidden similarities and differentiation among the samples. The internally validation of the PCA model was carried out using a leave one out full cross validation procedure, which involved investigating the maximum of 7 principle components.

### Pre-processing of NIR Spectral Data

Prior to PCA modeling, unit vector normalization, standard normal variate (SNV) and multiplicative scatter correction (MSC) were employed to correct the multiplicative and additive effects of the spectra. Smoothing is usually employed to eliminate instrumental clatter or background information and de-trending methods are usually implemented to reduce the effects of accumulating data sets from a trend. First and second derivatives of the spectra (D1, D2) based on the Savitzky–Golay algorithm with five smoothing points and polynomial order of 2, were also implemented to increase the spectral resolution. Derivatives are commonly used to reduce insignificant baseline signals from samples (Berthomieu and Hienerwadel, 2009).

### Light Microscopy

Tissue samples were fixed in 10% buffered formaldehyde solution for 8 h and then sliced into small pieces. The tissue pieces were dehydrated in graded concentrations of ethyl alcohol, diaphanized in xylol, impregnated and embedded in paraffin wax. Thin sections (4μm thickness) where produced by rotary microtome and then stained with Hematoxylin and Eosin (H&E). For morphological examination, slides of stained sections were screened using digital light microscope under magnifications ranging from 40X to 1000X.

## RESULTS

### FT-IR Analysis

FT-IR spectra represent the solid liver tissue samples in reflectance mode for all three groups in the wavenumber range of 4000–400 cm^-1^ (Fig. 1). FT-IR spectra demonstrated a prominent increase in the peak intensities in the wavenumber range from 950 to 1650 cm-1, and from 2009 to 3568 cm^-1^ as KBA doses were increased in relevance from group-1 to group-3 which indicated there was a clear effect of KBA solubility on treatments. The FTIR spectra of pure KBA gives characteristic peaks at 3437 cm1 (OH stretching), 2932 cm1 (C-H stretching), 1697 cm1 (C ¼ O stretching of aryl acid), 1453 cm1 (C-H bend), 1375 cm1 (COO symmetric stretching of carboxylates), 1240 cm1 (C-CO-C stretching of aryl ketone), 1025 cm1 and 988 cm1 (ring structures of cyclohexane) (Ruah et al., 2014).

**Fig. 1.**
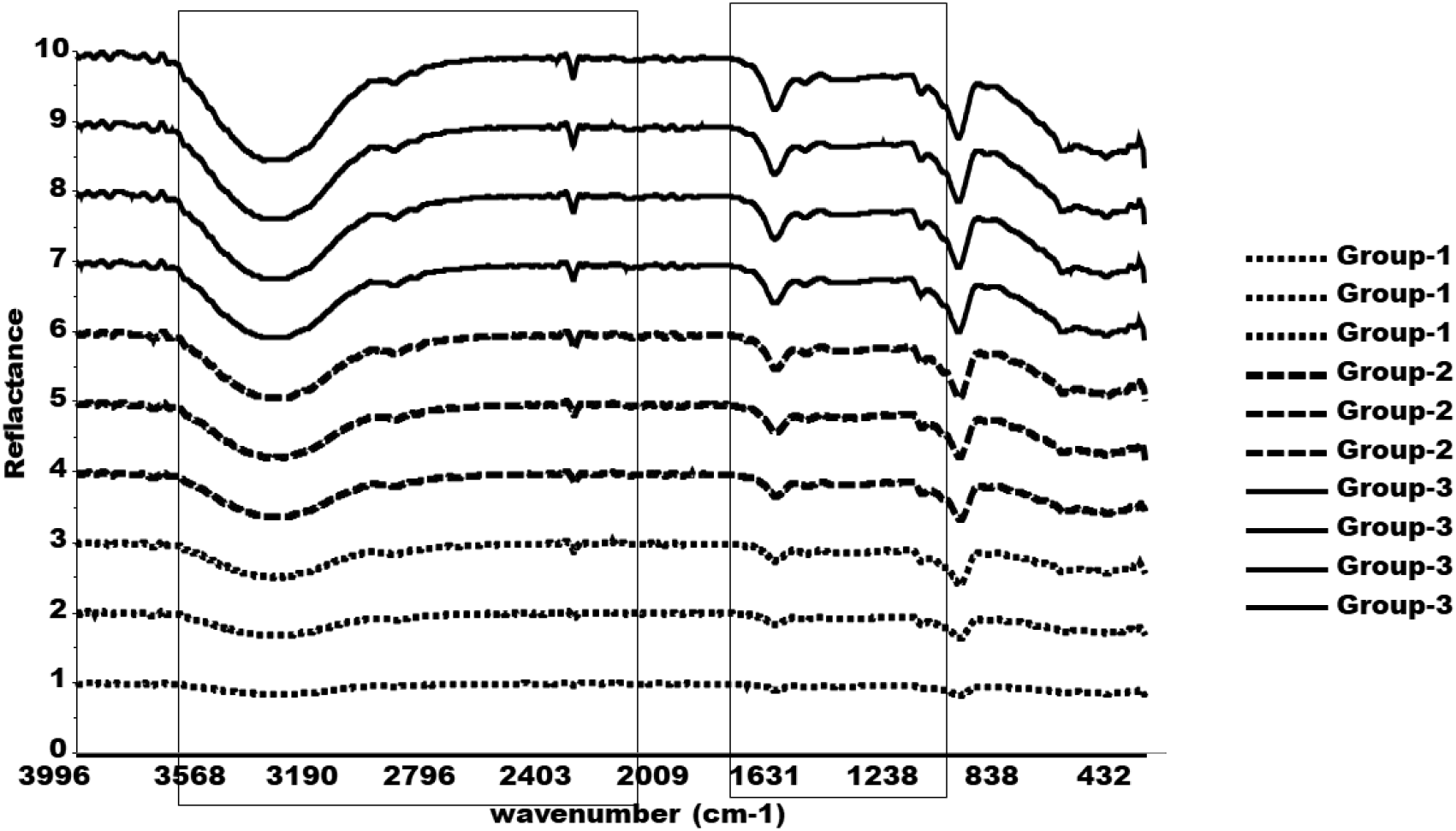
FT-IR spectra of solid liver tissue samples in reflectance mode for the three groups of mice. Group-1 normal liver tissue, group-2 diabetic control liver tissue, and group-3 treated IP with KBA tissue.

### Principle Component Analysis (PCA)

Principle Component Analysis (PCA) is a non-supervised multivariate exploratory data analysis tool to extract hidden information such as similarities and differences based on variations in the data. Therefore, PCA model (Fig. 2) was built to extract masked information of the FT-IR spectra for classification and discrimination among the three groups of liver tissue samples based on KBA solubility.

**Fig. 2.**
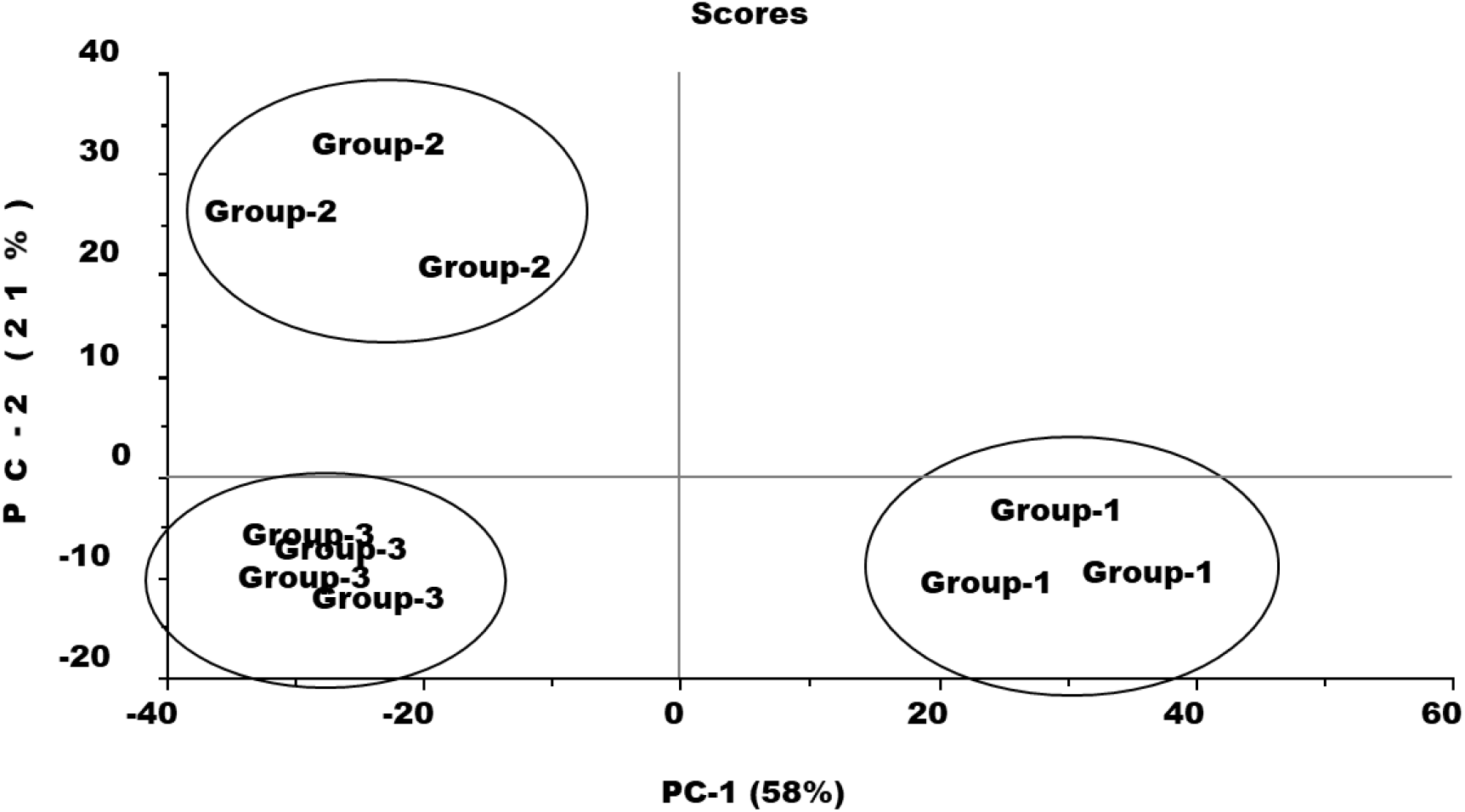
Principle Component Analysis (PCA) score plot for 3 groups of mice liver samples (group-1 normal liver, group-2 diabetic control liver, and group-3 diabetic liver treated IP with KBA).

The PCA score plot (Fig. 2) demonstrated a clear classification among the three groups of tissue samples based on the variation in FT-IR peak intensities as well as peak position. The three groups of liver tissue samples were mapped based on the solubility of the KBA. Samples of group-1 and group-3 are seperated on PC-1 axis that utilized 58% of spectral variation, while samples of group-2 and group-3 seperated on PC-2 axis carrying 21% of spectral information.

PCA loadings were plotted to determine the part of spectral variation contributed to the PCA model (Fig. 3). It was clear from the loading plots for PC-1 that the spectral regions in the wavenumber range from 950 to 1650 cm^-1^ and then from 2009 to 3568 cm^-1^ are the most import regions of the FT-IR spectra and contributed mainly to the PCA model in relevance from group-1 to group-3 liver tissue samples, indicated that there was a clear effect of KBA solubility on treatments as KBA concentrations peaks were distinctly elevated in group 3.

**Fig. 3.**
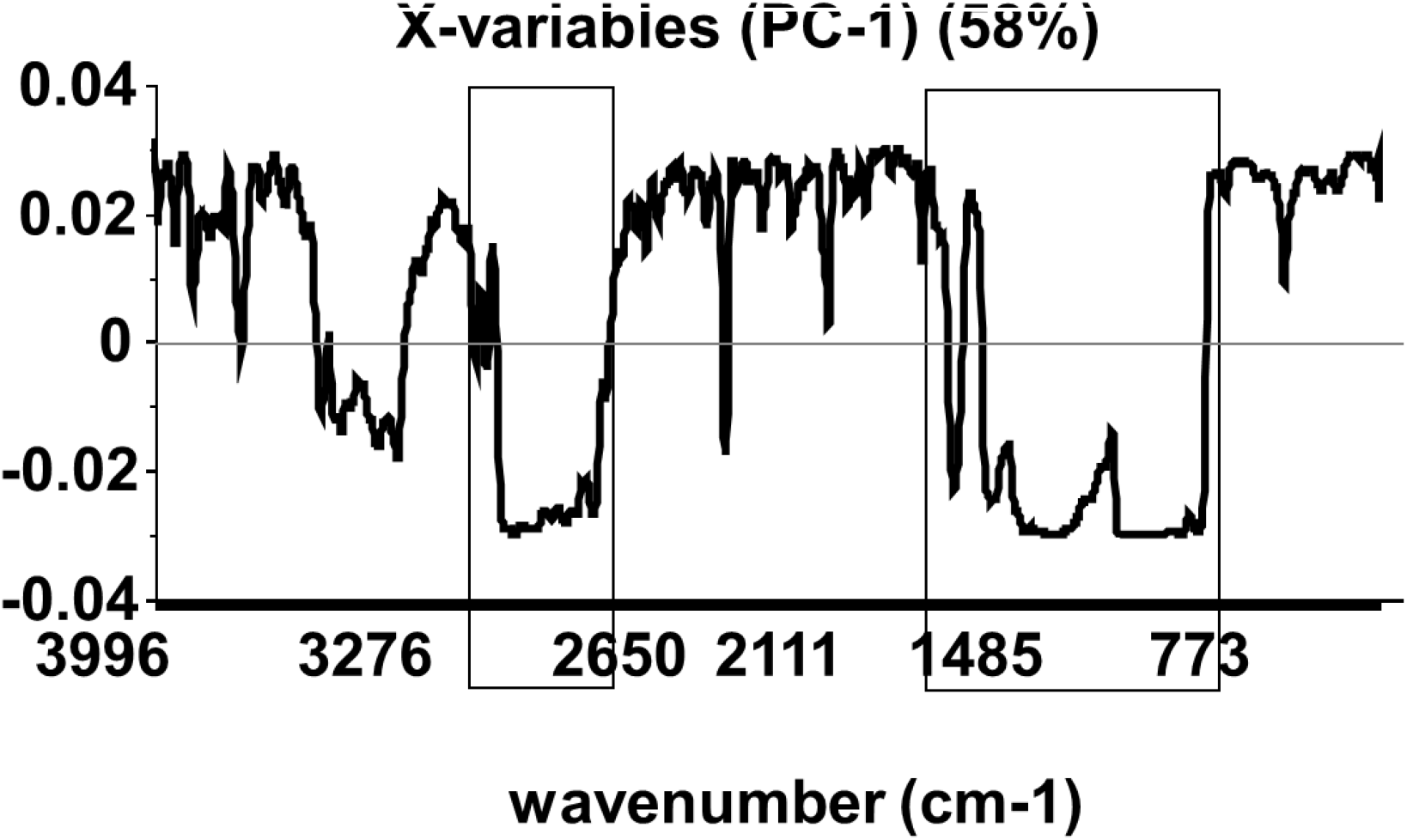
Principle Component Analysis (PCA) loading plot.

### Histopathological analysis

Liver sections of normal mice treated with citrate buffer (group 1) showed liver architecture made up of classical lobules and portal lobules (Fig 4A). The classical lobules consist of plates of hepatocytes radiating from a central vein and extending towards portal areas (Fig 4B). The portal lobules (triads) consist of hepatic artery, hepatic portal vein, bile duct and lymphatic vessels (Fig 4). Between the hepatocyte plates are liver sinusoids, which are capillaries that carry blood from the hepatic portal vein and hepatic artery enters via the portal trials, and then drains to the central vein (Figs 4B & 4C).

**Fig 4.**
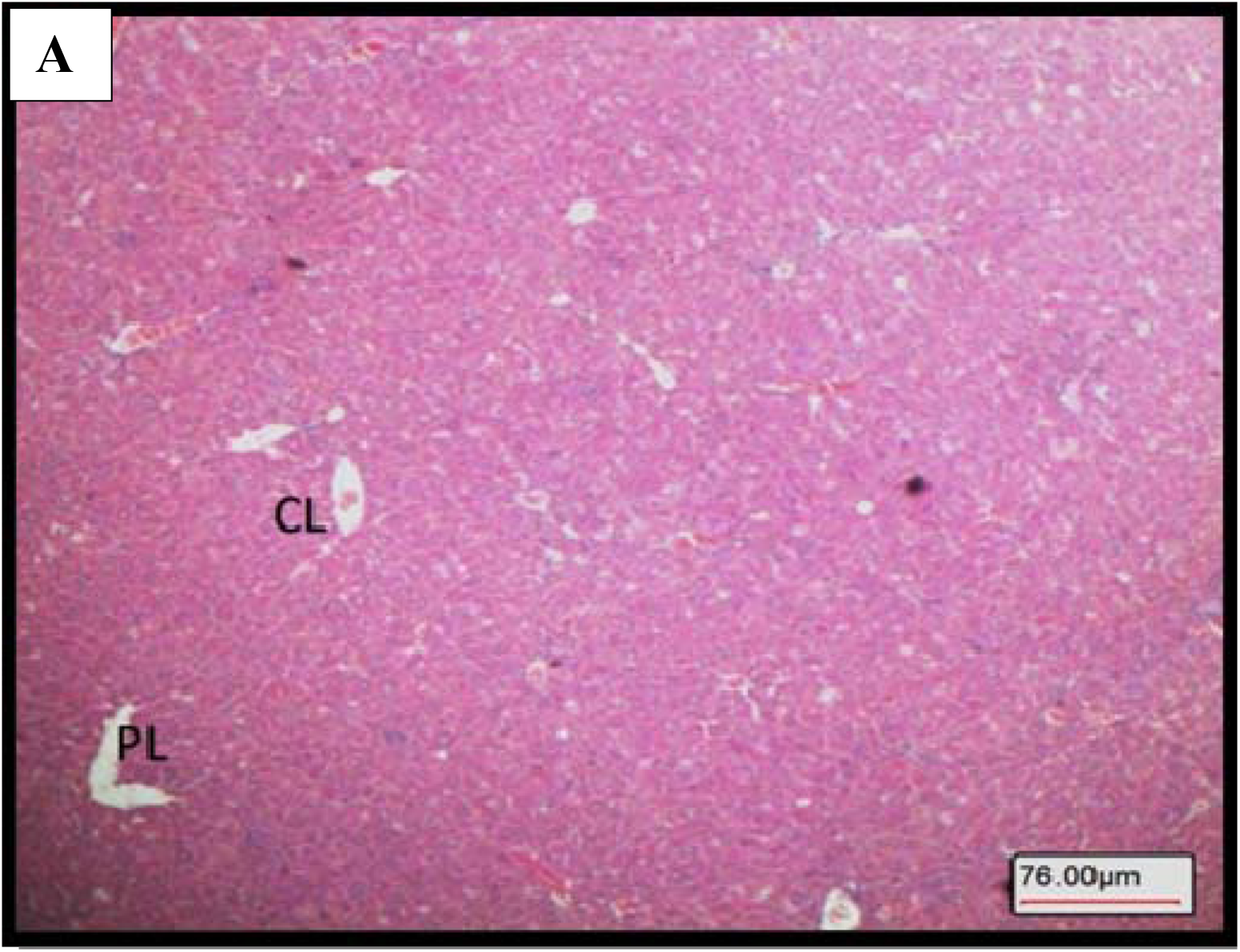
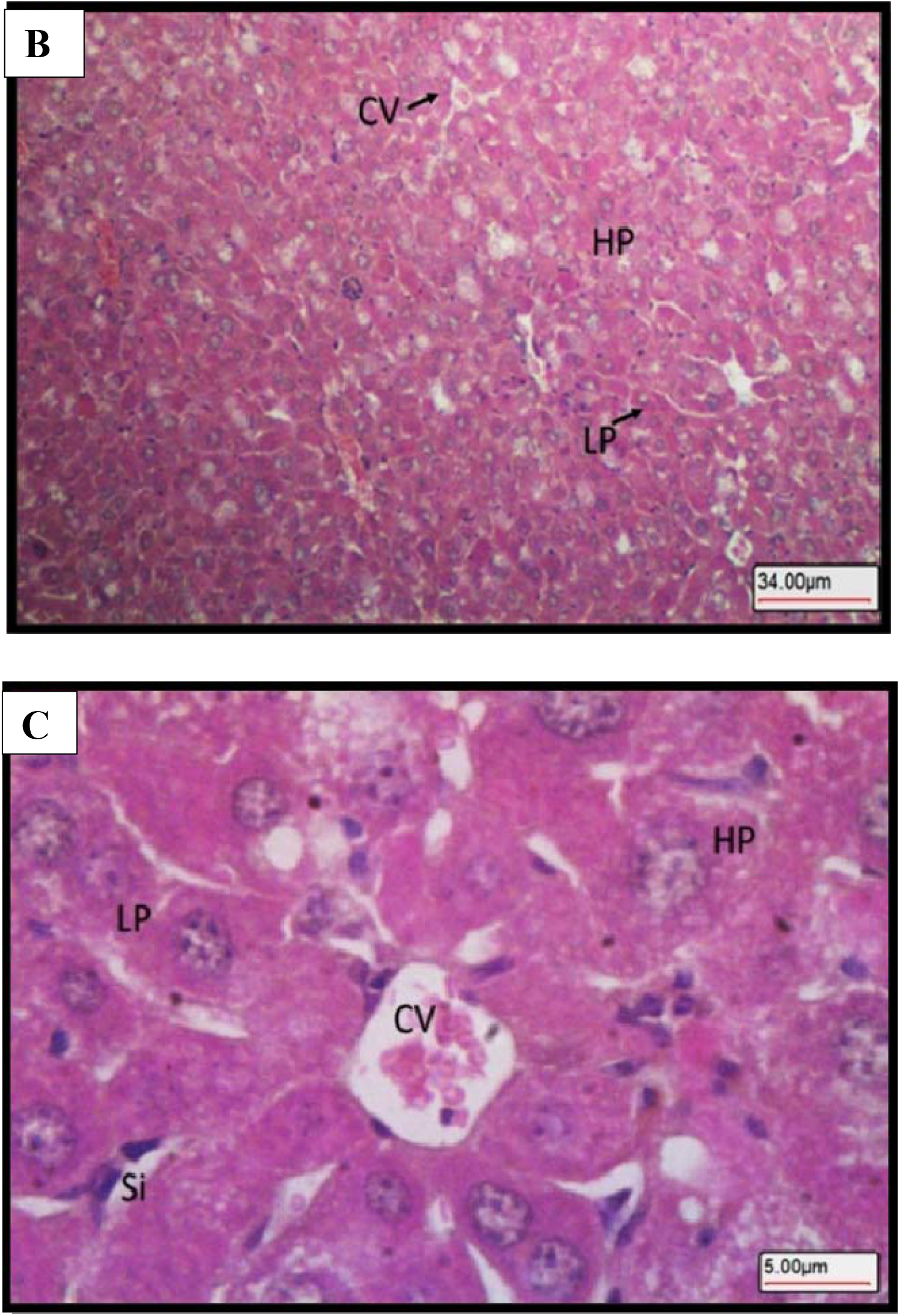
Light microscopy images of H&E-stained liver section showing normal control mice Group-1. A. Liver histology made up of classical lobules (CL) and portal lobules (PL). B. Classical lobules consist of plates of hepatocytes (HP) radiating from a central vein (CV). C. Central vein (CV) lined with epithelium (Ep), hepatocytes (H), liver hepatocyte plates (LP), liver sinusoids (Si). Mags. 40X, 100X. 600X, respectively.

Histological results of diabetic mice (group 2) showed that the classical and portal lobules of liver tissue (Fig 5A) with histopathological changes including hepatocellular damage in the form of increase vacuolation in the cytoplasm of hepatocytes appeared as indistinct clear vacuoles indicating glycogen infiltration and accumulation in diabetes (Figs 5B & 5C). Liver diabetic mice (group 3) treated IP with KBA showed normal liver architecture (Fig 6A) with mild vacuolations of hepatocytes (Figs 6B & 6C).

**Fig 5.**
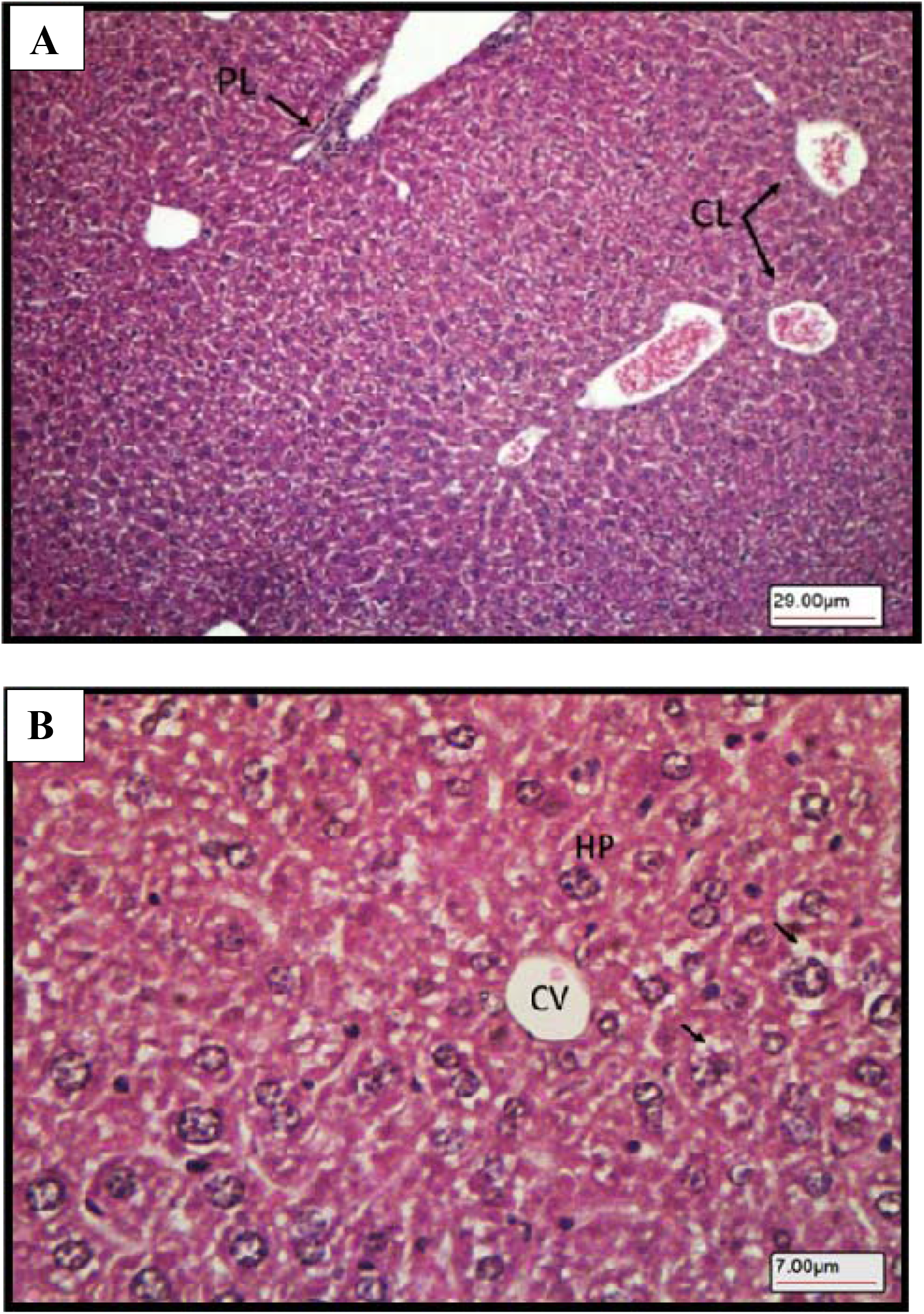
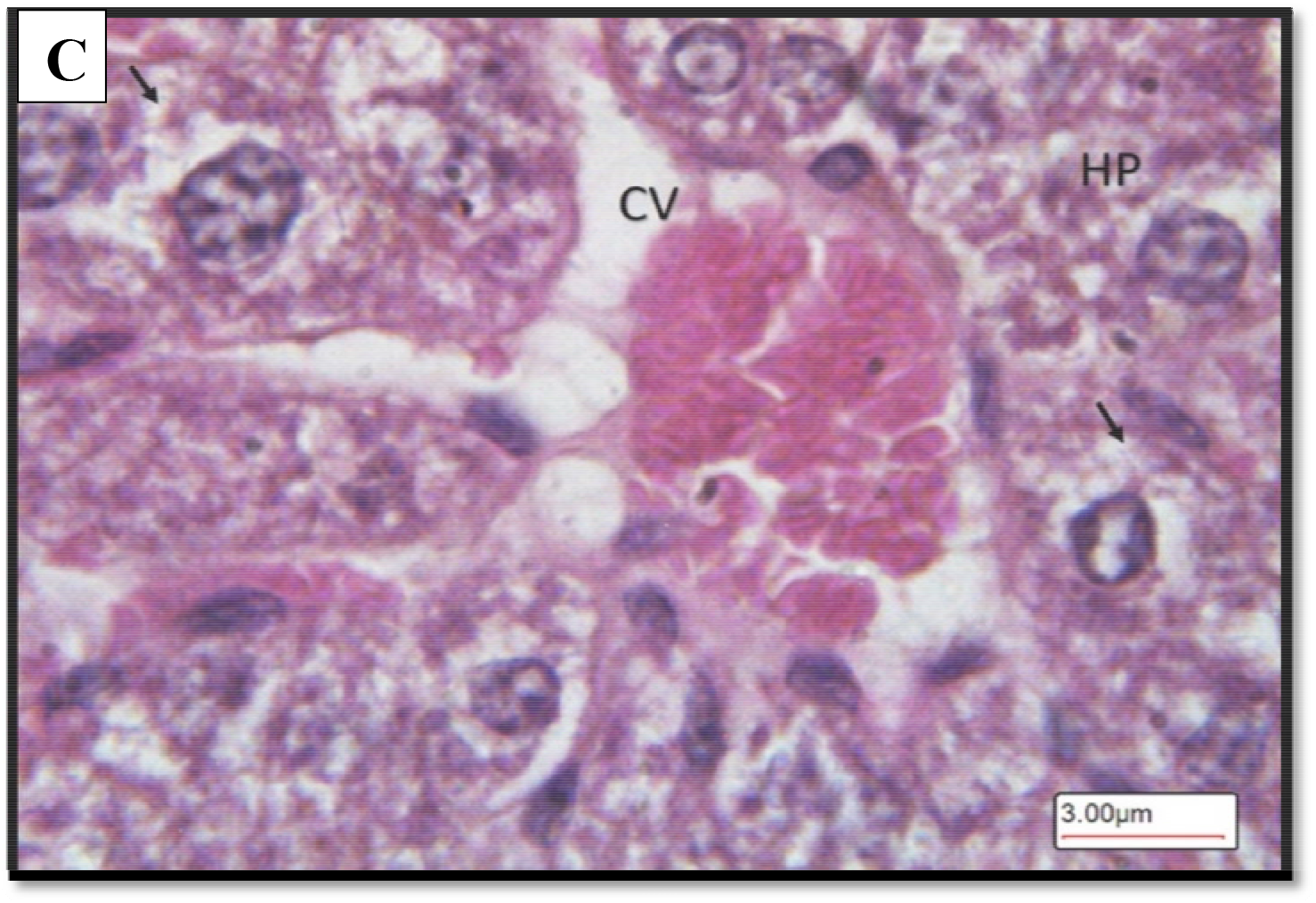
Diabetic control mice Group-2. A. Liver tissue architecture. B. hepatocellular damage in the form of increase vacuolation in the cytoplasm of hepatocytes appeared as indistinct clear vacuoles (arrows) indicate glycogen infiltration and accumulation in diabetes. Central vein (CV), Hepatocytes (HP). C. Higher magnification image showing vacuoles in the cytoplasm of hepatocytes (arrows). Central vein (CV), Hepatocytes (HP). Mags. 100X, 400X.,1000X, respectively.

**Fig. 6.**
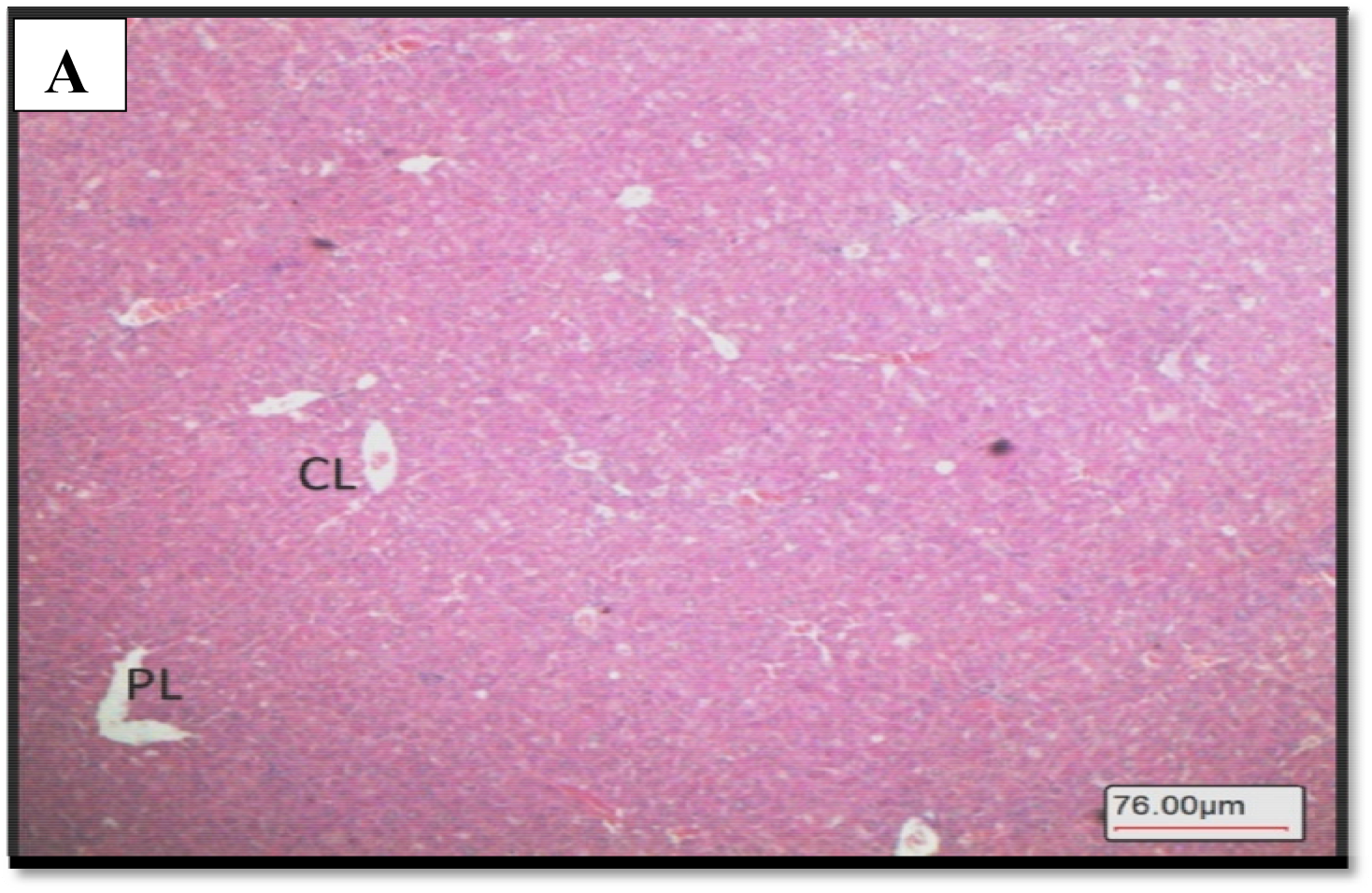
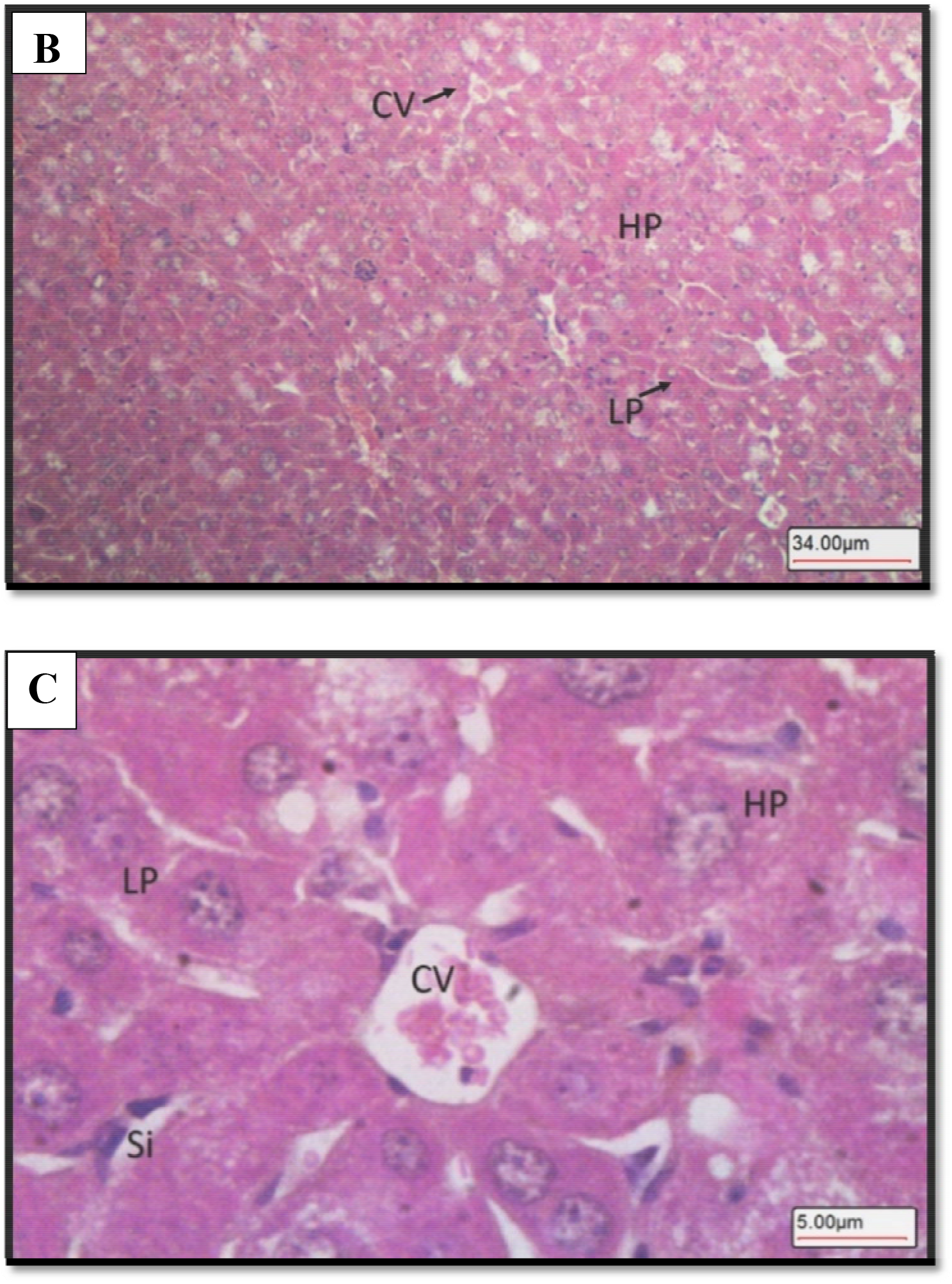
Diabetic mice IP treated with KBA Group-3. A. Liver tissue appeared with normal architecture of classical lobules (CL) and portal lobules (PL). B. Normal hepatocytes (HP) arranged in normal plates (LP) around central vein (CV). C. High magnification showing normal hepatocytes with mild vacuolations (HP) arranged in normal plates (LP) around central vein (CV). Liver sinusoids (Si). Mags. 40X, 400X, 600X, respectively.

## DISCUSSION

Diabetes mellitus is a metabolic disorder caused mainly by hyperglycemia and hyperlipidemia and affects most of the body’s vital organs and involves multiple therapies for treatment (Ginter and Simko, 2012). Currently, no efficient treatment available for certain types of diabetics (such as type 1 and late onset of autoimmune diabetes of adults) and recent studies reported that anti-diabetic properties of boswellic acids, the pharmacologically active constituents of frankincense can be used (Ammon, 2019). The anti-inflammatory properties of boswellic acids reported as targets for immune system factors involved in diabetes mellitus type 1 and type 2 (Ammon, 2016). However, boswellic acid compounds including α and β-boswellic acids (BA), and acetylated α and β-boswellic acids (ABA) showed no effects in inhibiting STZ-mediated hyperglycemia whereas 11-keto-β-boswellic acid (KBA), and 3-O-acetyl-11-keto-β-boswellic acid (AKBA), played an important role in the prevention of diabetes. These isolated compounds, KBA and AKBA, prevented the occurrence of autoimmune reactions, insulitis, and reduced hyperglycemia STZ-induced diabetes models (Shehata et al., 2015). Pharmacokinetic studies of *Boswellia* species extracts reported poor bioavailability, especially of KBA and KBA in both humans and rats owing to poor water solubility and high lipophilicity (Karlina et al., 2007; Krüger et al., 2008). Studies reported that permeability-associated barriers compromised oral bioavailability of KBA include its gastrointestinal volatility, mediated intestinal metabolism, accumulation within the enterocytes, and saturable kinetics (Bagul et al., 2014). In this study, a rapid and inexpensive detection FT-IR spectroscopy coupled with multivariate PCA method was developed to evaluate the solubility of KBA, from the gum resin extracted from the Omani frankincense in the liver of STZ induced diabetic mice. The newly developed method demonstrated that FT-IR-PCA was reliable and an accurate technique for the visualization of similarities and difference among the three groups of mice liver tissue samples based on the solubility of KBA and have indicated that there was a clear effect of treatments. The concentrations of KBA demonstrated distinct elevated peaks in FT-IR_PCA spectrum in the treated group. This showed that the concentration of KBA was improved following IP treatment and proved to be an effective route of administration leading to increased KBA bioavailability. Several studies used different approaches to enhance boswellic acids bioavailability (Skarke et al., 2012; Du et al., 2015) and many have reported in animal models of disease the biological effects of drugs after IP administration indicating bioavailability of these large molecules administered by this route (Lee et al., 2011; Zhang et al., 2014; Wangler et al., 2016; Chang et al., 2018). IP route of administration have proven to be more effective because the rate and extent of drug absorption is faster in IP followed by intramuscular than the oral routes (Wolfensohn and Lloyd, 1994). The IP administered pharmacological agents are exposed to a large surface area which leads to rapid and efficient absorption (Al Shoyaib et al., 2020). The present findings suggest high solubility of KBA in the animal model following IP treatment led to significant recovery of liver tissues and also evaluated the effect of KBA on the histological alterations in the liver of STZ induced diabetic mice. Streptozotocin (STZ) is a naturally occurring compound which selectively destroys beta cells and induces alkylation of DNA resulting in rapid necrosis, leading to hyperglycemia (Chinedum et al., 2013). Fresh liver tissue samples from the same groups used for the solubility test were also used for histological examination. The results showed a marked improvement in the liver tissues of diabetic mice with normal structures of hepatocytes exhibiting mild vacuolations in their cytoplasm. The results suggest that the active derivative of KBA from the gum resin of frankincense appeared to have a protective effect on the morphology of liver in diabetic mice by remarkable decreasing hypoglycemia caused by STZ. Several studies have confirmed the effectiveness of KBA for diabetes and suppressing the development of the disease in STZ-induced diabetic mice by triggering an immune response mediated by cytokines (Ammon, 2019; Shehata et al., 2015; Ammon et al. 2011). Furthermore, boswellic acids have been reported to minimize liver complications and provided a degree of protection (Iram et al., 2017). Liver tissue histological findings revealed that KBA has a protecting effect against diabetes-induced damage (Azemi et al., 2012). The results indicated that the current study used a reliable, rapid and inexpensive technique to support high separation among the different groups of mice liver tissues using FT-IR spectroscopy coupled with PCA based on solubility of KBA that significantly improved following IP treatment which led to significant recovery of the liver and hepatocytes.

Reflectance FT-IR spectroscopy coupled with PCA could be deployed for rapid, inexpensive and non-destructive detection method to examine the IP treatment effect of KBA solubility in mice liver tissue samples. Liver tissue histopathology revealed that KBA has a protecting effect against STZ induced-diabetes damage, which provided supporting evidence on the improvement of KBA absorption by liver tissue following IP treatment.

## ACKNOWLEDGMENTS

Laboratory for Stem Cells and Regenerative Medicine at Natural and Medical Sciences Research Centre and DARIS Research Centre, University, University of Nizwa which provided the facilities to carry out the project is highly appreciated. This work was supported by University of Nizwa, Sultanate of Oman. The use of the animals was approved by the Animal Ethics Committee at University of Nizwa.

## AUTHOR CONTRIBUTIONS

I.A, F.M, A.G, and S.A. conceived the experimental design. I.A. G.A., F.M, S.K., J.A, and J.S. performed the experiment, data observation and acquisition. I.A, F.M, I.K, A.A, A.H, S.H, and B.A analysed and interpreted the data. I.A, F.M and I.K, wrote the paper. All authors reviewed the manuscript, read and approved the final manuscript. The authors declare that they have no conflict of interest.

## REFERENCES

Ahmed HH, Ahmed A, Abd-Rabou AZ, Hassan S, Kotob E. (2015). Phytochemical Analysis and Anti-cancer Investigation of Boswellia Serrata Bioactive Constituents In Vitro. Asian Pacific Journal of Cancer Prevention, 16: 7179–7188. https://doi.10.7314/apjcp.2015.16.16.7179

Al-Haddad R, Karnib, N, Assaad RA, Bilen Y, Emmanuel N, Ghanem A, Younes J, Zibara V, Stephan JS, Sleiman SF. (2016). Epigenetic changes in diabetes. Neuroscience Letter, 625: 64–69. https://doi.10.1016/j.neulet.2016.04.046.

Al-Harrasi A, Ali L, Ceniviva E, Al-Rawahi A., Hussain, J., Hussain, H., Rehman, N.U., Abbas, G. and Al-Harrasi, R. (2013). Antiglycation and antioxidant activities and HPTLC analysis of Boswellia sacra Oleogum resin: the sacred frankincense. Tropical Journal of Pharmacological Research, 12: 597–602. https://doi.10.4314/tjpr.v12i4.23.

Al-Harrasi A, Ali L, Hussain J, Rehman NU, Ahmed M, Al-Rawahi A. (2014). Analgesic effects of crude extracts and fractions of Omani frankincense obtained from traditional medicinal plant Boswellia sacra on animal models. Asian Pacific Journal of Tropical Medicine, 7: S485–90. https://doi.10.1016/S1995-7645(14)60279-0.

Al-Shoyaib A, Archie SR, Karamyan VT. (2020). Intraperitoneal Route of Drug Administration: Should it Be Used in Experimental Animal Studies? Pharmacology Research, 37: 12, 1–17. https://doi.org/10.1007/si1095-019-2745-x.

Al-Yasiry AR, Kiczorowska B. (2016). Frankincense--therapeutic properties. Postępy Higieny i Medycyny Doswiadczalnej, 70: 380–91. https://doi.10.5604/17322693.1200553.

Ammon HPT. (2016). Boswellic acids and their role in chronic inflammatory diseases. Advances in Experimental Medicine and Biology, 928: 291–327. https://doi.10.1007/978-3-319-41334-1_13.

Ammon HP. (2019). Boswellic Extracts and 11-Keto-β-Boswellic Acids prevent Type 1 and Type 2 Diabetes Mellitus by suppressing the Expression of Proinflammatory Cytokines. Phytomedicine, 63; 153002. https://doi.10.1016/j.phymed.2019.153002.

Ammon H, Shehata AM, Quintanilla-Fend L. (2011). 11-Keto-beta-Boswellic Acid (KBA) Suppresses Development of Insulitis Apoptosis of Pancreatic Islets and Elevation of Proinflammatory Interleukins in Multiple Low Dose Streptozotocin (MLD-STZ)-Induced Diabetes of Mice. American Diabetes Association-in Diabetes Pro, 60: A278–A278.

Azemi ME, Namjoyan F, Khodayar MJ, Ahmadpour F, Padok AD, Panahim M. (2012). The antioxidant capacity and anti-diabetic effect of Boswellia serrata triana and planch aqueous extract in fertile female diabetic rats and the possible effects on reproduction and histological changes in the liver and kidneys. Journal of Natural Pharmaceutical Products, 7: 168–175. https://PMCID:PMC3941871

Bagul P, Khomane KS, Bansal AK. (2014). Investigating permeability related hurdles in oral delivery of 11-keto-beta-boswellic acid. International Journal of Pharmaceutics, 464: 104–110. https://doi.10.1016/j.ijpharm.2014.01.019.

Baynes HW. (2015). Classification, Pathophysiology, Diagnosis and Management of Diabetes Mellitus. Journal of Diabetes and Metabolic Disorders, 6: 1–9. https://doi.10.4172/2155-6156.1000541.

Berthomieu C, Hienerwadel R. (2009). Fourier transform infrared (FTIR) spectroscopy. Photosynthesis Research, 101: 157–70. https://doi.10.1007/s11120-009-9439-x

Büchele B, Simmet T. (2003). Analysis of 12 different pentacyclic triterpenic acids from frankincense in human plasma by high performance liquid chromatography and photodiode array detection. Journal of Chromatography, B 795: 355–62. https://doi.10.1016/s1570-0232(03)00555-5.

Chang R, Al-Maghribi A, Vanderpoel V, Vasilevko V, Cribbs DH, Boado R, Pardridge WM, Sumbria RK. (2018). Brain penetrating Bifunctional erythropoietin-transferrin receptor antibody fusion protein for Alzheimer’s disease. Molecular Pharmaceutics, 15: 4963–73. https://doi.10.1021/acs.molpharmaceut.8b00594.

Chinedum OE, Eleazu K, Chukwuma S, Essien UN. (2013). Review of the mechanism of cell death resulting from streptozotocin challenge in experimental animals, its practical use and potential risk to humans. Journal of Diabetes and Metabolic Disorders, 12:1–7. https://doi.10.1186/2251-6581-12-60

Cotoi CG, Quaglia A. (2016). Normal Liver Anatomy and Introduction to Liver Histology. In Textbook of Pediatric Gastroenterology, Hepatology and Nutrition. pp. 609–612. Springer, Cham.

Du Z, Liu Z, Ning Z, Liu Y, Song Z, Wang C, Lu A. (2015). Prospects of boswellic acids as potential pharmaceutics. Planta Medica, 81: 259–271. https://doi.10.1055/s-0034-1396313.

International Diabetic Federation (IDF). Idf.org. 2020 [cited Dec 2020]. Available from: https://idf.org/our-network/regions-members/middle-east-and-north-africa/members/42-oman.html.

Ginter E, Simko V. (2012). Type 2 Diabetes Mellitus, Pandemic in 21st Century. Advances in Experimental Medicine and Biology, 771: 42–50. doi: 10.1007/978-1-4614-5441-0-6.

Hüsch J, Bohnet J, Fricker G, Skarke C, Artaria C, Appendino G, Schubert-Zsilavecz M, Abdel-Tawab M. (2013). Enhanced absorption of boswellic acids by a lecithin delivery form (Phytosome®) of Boswellia extract. Fitoterapia, 84: 89–98. https://doi.10.1016/j.fitote.2012.10.002.

Iram F, Khan SA, Husain A. (2017). Phytochemistry and potential therapeutic actions of Boswellic acids: A mini-review. Asian Pacific Journal; of Tropical Biomedicine, 7: 513–23. https://doi.org/10.1016/j.apjtb.2017.05.001.

Jianbiao Yue, Yang X, Xia L. (2019). Wound Healing Potential of the Standardized Extract of Boswellia serrata on Experimental Diabetic Foot Ulcer via Inhibition of Inflammatory, Angiogenetic and Apoptotic Markers. Planta Medica, 85: 657–669. https://doi.10.1055/a-0881-3000.

Karlina MV, Pozharitskaya ON, Kosman VM, Ivanova SA. (2007). Bioavailability of boswellic acids: in vitro/in vivo correlation. Pharmaceutical Chemistry Journal, 41: 569–572. https://doi.10.1007/s11094-008-0017-x

Krüger P, Rambod D, Gunter P, Eckert JK, Dietrich A, Volmer UB, Walter E, Muller MK, Schubert-Zsilavecz M, Abdel-Tawab M. (2008). Metabolism of Boswellic Acids in Vitro and in Vivo. Drug Metabolism and Disposition, 36: 1135–1142. https://doi.org/10.1124/dmd.107.018424

Lee B, Clarke D, Al-Ahmad A, Kahle M, Parham C, Auckland L, Shaw C, Fidanboylu N, Orr WA, Ogunshola O, Fertala A, Thomas SA, Gregory JB. (2001). Perlecan domain V is neuroprotective and proangiogenic following ischemic stroke in rodents. Journal of Clinical Investigation, 121: 3005–23. https://doi.10.1083/jcb.201807178.

Mohamed J, Nafizah AN, Zariyantey AH, Budin SB. (2016). Mechanisms of diabetes-induced liver damage: the role of oxidative stress and inflammation. SQU Medical Journal, 16: e132. https://doi.10.18295/squmj.2016.16.02.002.

Middha SK, Usha T, Pande V. (2014). Letter to the editor: pomegranate peel atten PMCID: PMC4464388uates hyperglycemic effects of alloxan-induced diabetic rats. EXCLI Journal, 13: 223–4. https://PMC4464388

Ni X, Suhail MM, Yang Q, Cao A, Fung KM, Postier RG, Woolley C, Young G, Zhang J, Lin HK. (2012). Frankincense essential oil prepared from hydrodistillation of Boswellia sacra gum resins induces human pancreatic cancer cell death in cultures and in a xenograft murine model. BMC Compl. Alternative Medicine, 12: 253 (1-14). https://doi.10.1186/1472-6882-12-253.

Pengzong Z, Yuanmin L, Xiaoming X, Shang D, Wei X, Zhigang L, Dongzhou D, Wenjing Y, Jianbiao Y, Yang X, Xia L. (2019). Wound Healing Potential of the Standardized Extract of Boswellia serrata on Experimental Diabetic Foot Ulcer via Inhibition of Inflammatory, Angiogenetic and Apoptotic Markers. Planta Medica, 85: 657–669. https://doi.10.1080/21691401.2019.1699814.

Shehata A, Quintanilla-Fend L, Bettio S, Kamyabi-Moghaddam Z, Kohlhofer U, Scherbaum W, Ammon H. (2017). 11-Keto-β-Boswellic Acid Inhibits Lymphocyte (CD3) Infiltration into Pancreatic Islets of Young None Obese Diabetic (NOD) Mice. Hormone and Metabolic Research, 49: 693–700.

Sabra SM, Al-Masoudi LM. (2014). The effect of using frankincense (Boswellia sacra) chewing gum on the microbial contents of buccal/oral cavity, Taif, KSA. Journal of Medical and Dental Sciences, 13: 77–82. https://PMC3304380

Shaikh H, Shrivastava VK. (2014). Effects of steptozotocin induced diabetes mellitus type 1 on the rat brain antioxidant status and activity of acetylcholinesterase: a novel and potential treatment by Vitex negundo. International Journal of Pharmacy and Pharmaceutical Sciences, 6: 252–6. https://innovareacademics.in/journals/index.php/ijpps/article/view/2714.

Saraswati S, Agrawal SS. (2012). Antiangiogenic and cytotoxic activity of boswellic acid on breast cancer MCF-7 cells. Biomedicine and Preventive Nutrition, 2: 31–37.2012. https://doi10.1016/j.bionut.2011.09.006.

Sharma S, Thawani V, Hingorani M, Shrivastava V, Bhate R, Khiyani R. (2004). Pharmacokinetic study of 11Keto β-Boswellic Acid. Phytomedicine, 11: 255–260. https://doi.10.1078/0944-7113-00290.

Skarke C, Kuczka K, Tausch L, Werz O, Rossmanith T, Barrett JS, Harder S, Holtmeier W, Schwarz JA. (2012). Increased bioavailability of 11-keto-beta-boswellic acid following single oral dose frankincense extract administration after a standardized meal in healthy male volunteers: Modeling and simulation considerations for evaluating drug exposures. Journal of Clinical Pharmacology, 52: 1592–1600. https://doi.10.1177/0091270011422811.

Shehata A, Quintanilla-Fend L, Bettio S, Jauch J, Scior T, Scherbaum W, Ammon H. (2015). 11-Keto-β-Boswellic Acids Prevent Development of Autoimmune Reactions, Insulitis and Reduce Hyperglycemia During Induction of Multiple Low-Dose Streptozotocin (MLD-STZ) Diabetes in Mice. Hormone and Metabolic Research, 47: 463–469. https://doi.10.1055/s-0035-1547293

Targher G, Lonardo A, Byrne CD. (2018). Nonalcoholic fatty liver disease and chronic vascular complications of diabetes mellitus. Nature Reviews Endocrinology, 14: 99. https://doi.10.1038/nrendo.2017.173.

Ruah ME, Rasaruddin NF, Fong SS, Jaafar MZ. (2014). Data preprocessing methods of FT-NIR spectral data for the classification cooking oil. American Institute of Physics Conference Proceedings, 1635: 890–897. https://doi.org/10.1063/1.4903688.

Wang H, Zhang C, Wu Y, Ai Y, Lee DY, Dai R. (2014). Comparative pharmacokinetic study of two boswellic acids in normal and arthritic rat plasma after oral administration of Boswellia serrata extract or Huo Luo Xiao Ling Dan by LC□MS. Biomedical Chromatography, 28:1402–1408. https://doi.10.1002/bmc.3182.

Wangler NJ, Jayaraman S, Zhu R, Mechref Y, Abbruscato TJ, Bickel U, Karamyan V. 2016. Preparation and preliminary characterization of recombinant neurolysin for in vivo studies. Journal of Biotechnology, 234: 105–15. https://doi.10.1016/j.jbiotec.2016.07.007.

Wolfensohn SE, Lloyd MH. (1994). Aleutian disease in laboratory ferrets. Veterinary Record, 134: 100. https://doi.10.1136/vr.134.4.100

Zhang B, Dai J, Wang H, Wei H, Zhao J, Guo Y, Fan K. (2014). Antiosteopontin monoclonal antibody prevents ovariectomy-induced osteoporosis in mice by promotion of osteoclast apoptosis. Biochemical and Biophysical Research Communications, 452: 795–800. https://doi.10.1016/j.bbrc.2014.08.149.

